# Expansion of Human Papillomavirus-Specific T Cells in Periphery and Cervix in a Therapeutic Vaccine Recipient Whose Cervical High-Grade Squamous Intraepithelial Lesion Regressed

**DOI:** 10.1101/2020.11.07.364844

**Authors:** Takeo Shibata, Sumit Shah, Teresa Evans, Hannah Coleman, Benjamin J. Lieblong, Horace J. Spencer, C. Matthew Quick, Toshiyuki Sasagawa, Owen W. Stephens, Erich Peterson, Donald Johann, Yong-Chen Lu, Mayumi Nakagawa

## Abstract

Advances in high-throughput sequencing have revolutionized the manner with which we can study T cell responses. We describe a woman who received a human papillomavirus (HPV) therapeutic vaccine called PepCan, and experienced complete resolution of her cervical high-grade squamous intraepithelial lesion. By performing bulk T cell receptor (TCR) β deep sequencing of peripheral blood mononuclear cells before and after 4 vaccinations, 70 putatively vaccine-specific clonotypes were identified for being significantly increased using a beta-binomial model. Interferon-γ enzyme-linked immunospot assay identified significantly increased HPV-specific T cell response in the HPV 16 E6 91-115 region after 4 vaccinations (*p*=0.023). T cells with specificity to this region were sorted and analyzed using single-cell RNA-seq and TCR sequencing, HPV specificity in 60 of the 70 clonotypes identified to be vaccine-specific was demonstrated. TCR β bulk sequencing of the cervical liquid-based cytology samples and cervical formalin-fixed paraffin-embedded samples before and after 4 vaccinations demonstrated the presence of these HPV-specific T cells in the cervix. Combining traditional and cutting-edge immunomonitoring techniques enabled us to demonstrate expansion of HPV-antigen specific T cells in the periphery and cervix.

## Introduction

Human papillomavirus (HPV) is best known as the causative agent of cervical cancer, but it can also cause cancers at other mucosal sites including the anus, oropharynx, penis, vagina, and vulva. It is estimated that HPV is responsible for 42,700 cancers in the US each year,^1^ including more than 90% of anal and cervical cancers and about 70% of oropharyngeal, vaginal, and vulvar cancers.^1^ Incidences of HPV-associated anal and oropharyngeal cancers have increased notably, and, although incidence of cervical cancer has stabilized after significant decreases over the past several decades,^2^ this remains the fourth most common cancer among women globally.^3^ The available prophylactic vaccines are effective for preventing HPV infections, but they cannot eliminate established infections; therapeutic vaccines could fill this need, but none are currently available.^4^ Such vaccines would benefit young women (narrowly, those ≤24 years old) and, broadly, any woman who plans to become pregnant^5^ because increased incidence of preterm delivery (from 4.4% to 8.9%) is associated with surgical treatments (e.g., loop electrical excision procedure [LEEP]) for high-grade squamous intraepithelial lesion (HSIL).^5,6^

We evaluated the safety of an HPV therapeutic vaccine (PepCan) in a single-center, single-arm, dose-escalation Phase I clinical trial treating women with biopsy-proven HSILs (NCT01653249).^7,8^ PepCan consists of four current good manufacturing practice (cGMP)-grade peptides covering the human papillomavirus type 16 (HPV 16) E6 protein (amino acids 1-45, 46-80, 81-115, and 116-158) and *Candida albicans* skin test reagent (Candin®, Nielsen Biosciences, San Diego, CA). PepCan was shown to be safe, and resulted in a histological regression rate of 45% which is roughly double that of a historical placebo (22%).^9^ In addition, circulating, peripheral T-helper type 1 (Th1) cells (*p*=0.0004) was increased, and the HPV 16 viral load was significantly decreased (*p*=0.008).^7^

Recent advances in high-throughput sequencing technology have enhanced our ability to appreciate how the T cell receptor (TCR) repertoire may reveal the role of T cells in immunotherapy for HPV-related diseases.^10–12^ The actual diversity present in a human body is estimated to be around 10^13^ unique TCRs.^13^ Next generation sequencing can facilitate the simultaneous analysis of millions of TCR sequences. Understanding the cytotoxic T cell repertoire, in parallel with observing clinical responses, would be essential for revealing immune mechanisms behind immunotherapies for chronic infectious diseases or cancer.^10,14–17^ In this article, we utilize multiplexed PCR-based TCR sequencing using genomic DNA to characterize TCR repertoires in peripheral blood mononuclear cells (PBMCs), stimulated CD3 cells, formalin-fixed paraffin-embedded (FFPE) tissues, and liquid-based cytology (LBC) samples from one subject who was a histologic responder from the Phase I clinical trial mentioned above. In addition, single-cell RNA-seq and TCR sequencing approaches were utilized to reveal the TCR sequences of HPV-specific T cells with a specificity to the HPV 16 91-115 amino acid region revealed by the enzyme-linked immunospot (ELISPOT) assay. We provide a proof-of-principle that a traditional immune assay, such as ELISPOT, can be combined with a cutting-edge technology to characterize HPV-specific T cells.

## Results

### Clinical trial design and vaccine response

The subject, a 41-year old Caucasian woman, participated in a single-arm, open-label Phase I clinical trial of an HPV therapeutic vaccine, PepCan, for treating biopsy-proven cervical HSILs (Figure 1A).^7,8^ At study entry, she had cervical intraepithelial neoplasia 3 (CIN 3), and was positive for HPV types 16, 31, and 58. At study exit (12 weeks after vaccination series completion), her LEEP biopsy was benign but was noted to have marked lymphocytic infiltration. Furthermore, she was noted to have leukocytosis and lymphocytosis (Table 1), and was positive for HPV 40 at exit. ELISPOT assay showed pre-existing HPV-specific response and responses to multiple HPV regions detected after vaccination. The response to one region, HPV 16 E6 91-115, was significantly increased (Figure 1B, *p*=0.023). Peripheral immune cell profiling showed increased percentage of Th1 cells, but unchanged levels of Tregs and Th2 cells (Figure 1C). Her HLA types were HLA-A*24/A*30, B*15/B*51, C*01/C*03, DPB1*02/ DPB1*02, DQB1*03/DQB1*06, and DRB1*11/DRB1*13.

**Table 1.** Complete blood count with differentials.

| Test | Reference range | Pre | Post-2 | Post-4 |
| --- | --- | --- | --- | --- |
| WBC (K/μL) | 3-12 | 7.92 | 8.14 | **13.94** |
| Hemoglobin (g/dL) | 11.5-16 | 12.9 | 13.4 | 13.9 |
| Hematocrit (%) | 34-47 | 39.1 | 41.2 | 42.7 |
| Platelet (K/μL) | 150-500 | 225 | 225 | 237 |
| Neutrophils (K/μL) | 2.5-8.2 | 4.5 | 5 | 7 |
| Lymphocyte (K/μL) | 1-4.8 | 2.5 | 2.3 | **5.5** |
| Monocytes (K/μL) | 0.1-1 | 0.6 | 0.6 | 0.9 |
| Eosinophils (K/μL) | 0-0.4 | 0.3 | 0.2 | 0.4 |
| Basophils (K/μL) | 0-0.22 | 0.02 | 0.02 | 0.03 |

Bold indicates values outside of the reference range.

**Figure 1.**
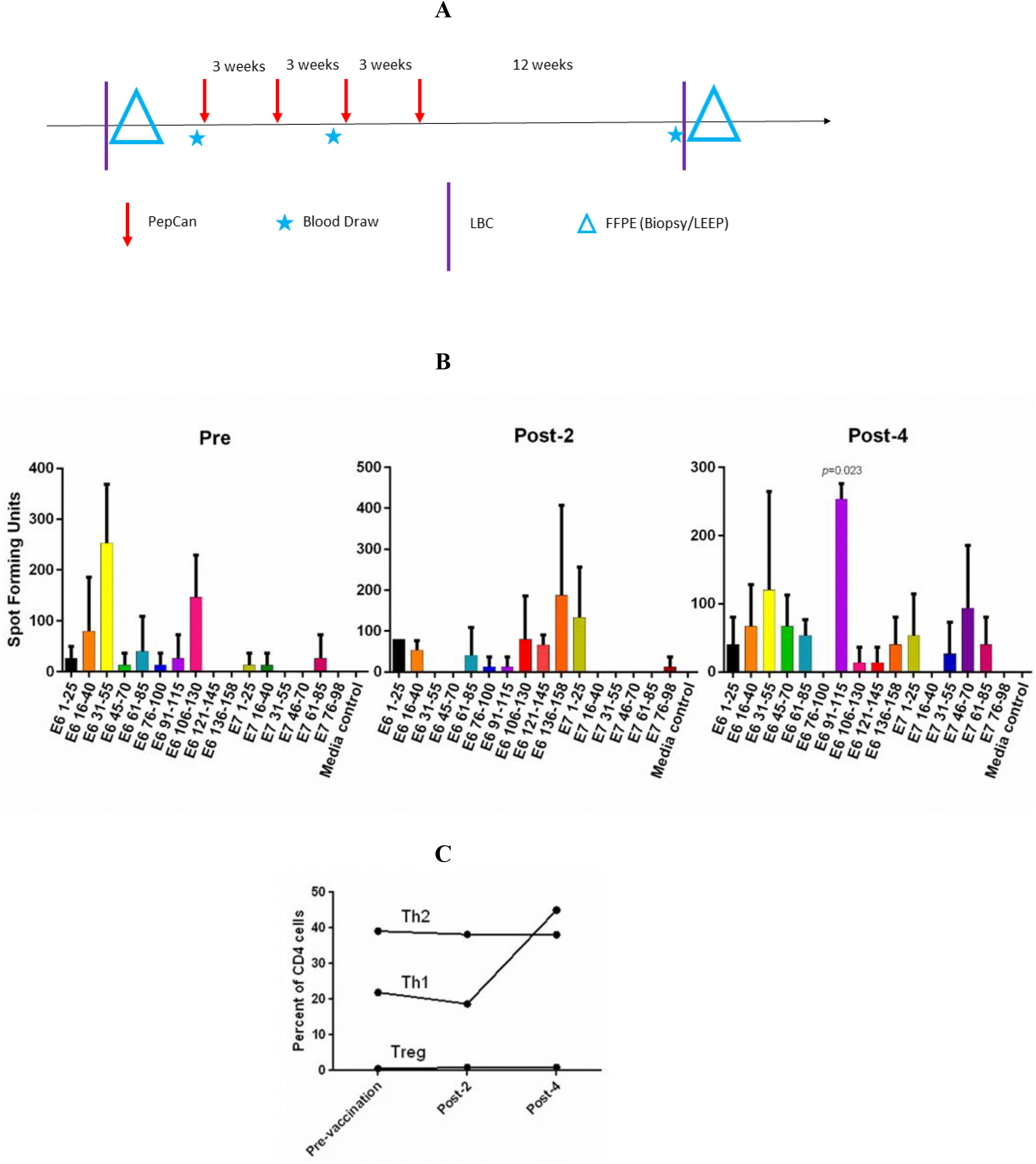
The Phase I clinical trial design and routine immune monitoring assays. **A.** Clinical trial design of the Phase I study. Vaccination (PepCan) visits were scheduled 3 weeks apart for patients who had biopsy-confirmed cervical high-grade squamous intraepithelial lesions (HSILs, i.e. CIN grade 2 or 3). Blood draws were performed pre-vaccination, and post-2 and post-4 vaccinations. Cervical local samples (LBC and FFPE) were collected pre-vaccination and post-4 vaccinations. FFPE samples were prepared from a pre-vaccination cervical biopsy and from loop electrical excision procedure (LEEP) biopsy post-4 vaccinations. **B.** Immunogenic HPV16 E6 and E7 regions were determined for each vaccine phase using ELISPOT assay. In pre-vaccine phase, positive responses (i.e., at least twice the media control) were detected in the E6 16-40, E6 31-55, and E6 106-130 regions. Positive responses were seen in the E6 1-25, E6 106-130, E6 136-158, and E7 1-25 regions in the post-2 vaccination sample, and in the E6 31-55, E6 91-115, and E7 46-70 regions in the post-4 vaccination sample. The increase in the response to the HPV16 E6 91-115 regions was statistically significant (paired *t*-test, *p*=0.023). Phytohemagglutinin was used as a positive control (not shown). The bars represent mean spot forming units per 1 × 10^6^ CD3 T cells, and error bars represent standard error of means. **C.** The Th1 level expressed as the percentage of CD4 T cells increased after 4 vaccinations, but Treg and Th2 levels were minimally changed.

### Multiplexed PCR-based TCR β chain deep sequencing

All samples examined (n=10: PBMC and stimulated CD3 samples at pre-, post-2, and post-4 vaccinations; and FFPE and LBC samples at pre- and post-4 vaccinations) yielded sufficient quantities of DNA for bulk TCR sequencing. In total, 749,417 clonotypes, and 1,256,277 T cells were identified in these 10 samples (Table 2). The numbers of total T cells and clonotypes were higher in PBMC than in stimulated CD3 (Figure 2). In cervical samples, the clonotypes (372,344,699 and 6,748) and total T cells (403,382,814, and 10,731) were detected in FFPE (pre and post-4) and LBC (pre and post-4). The productive clonality was increased after 4 vaccinations in PBMC, stimulated CD3, and LBC samples, and the maximum productive frequencies at least doubled in all sample types (Figure 2). The T cell fraction was highest in stimulated CD3 cells, and lowest in LBCs. DNA per cell was similar among PBMC, stimulated CD3 cells, and LBC (ranging from 0.0061 ng/cell to 0.011 ng/cell), but much higher in FFPE samples (0.714 ng/cell for pre-vaccination and 1.27 ng/cell for post-4 vaccinations), possibly reflecting lower quality DNA.

**Table 2.**
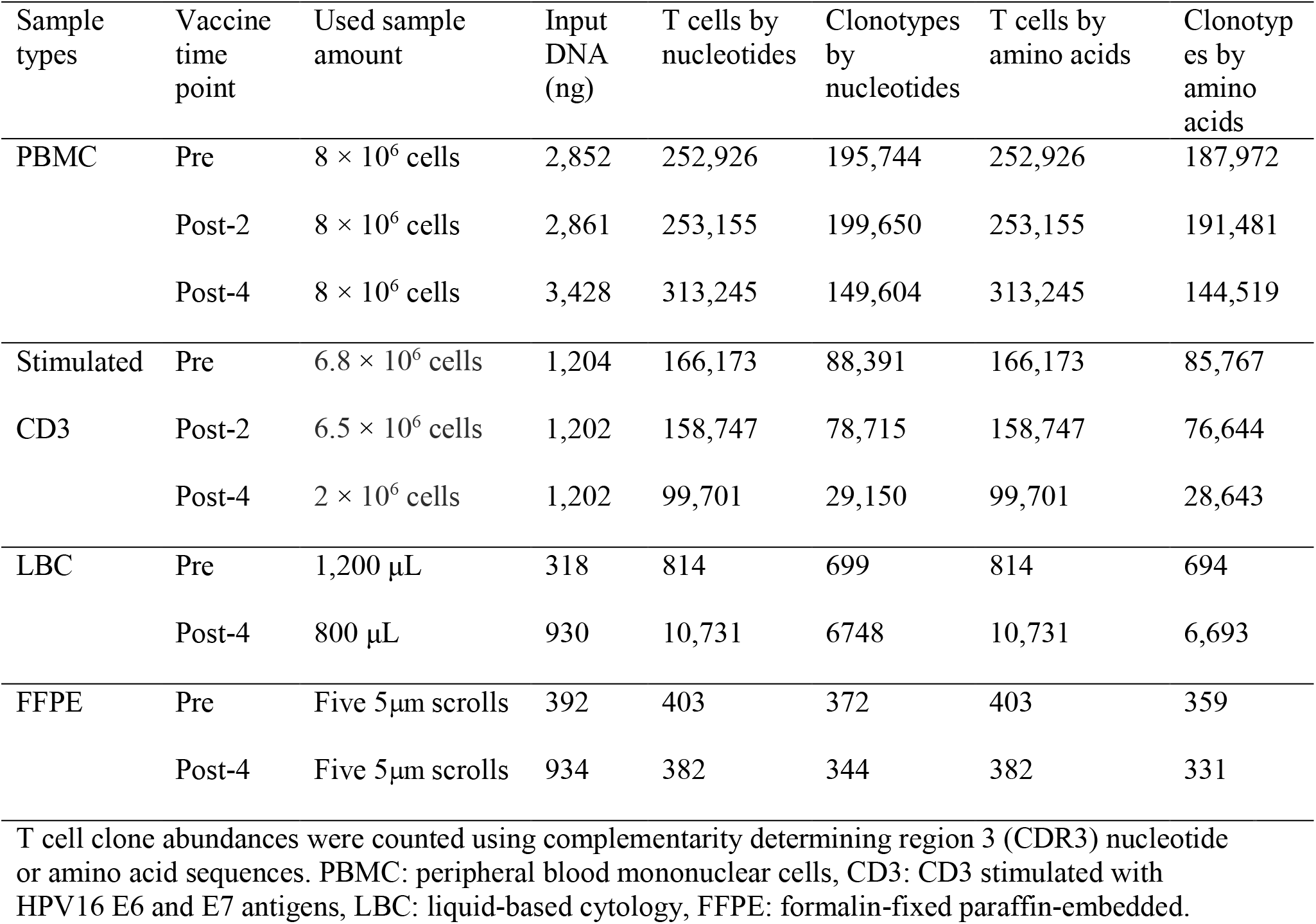
Sample characteristics.

**Figure 2.**
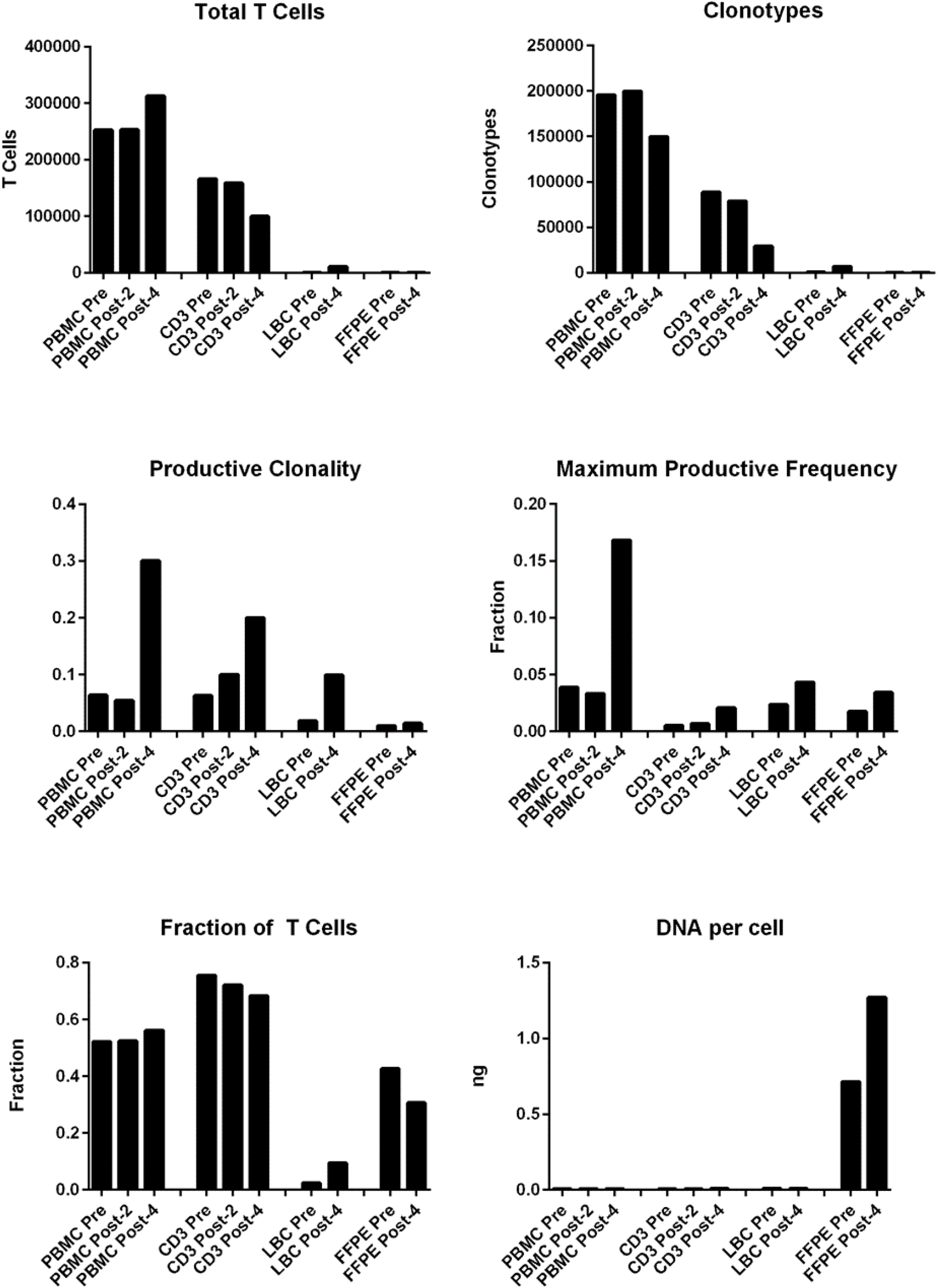
T cell structures of PBMC stimulated CD3, LBC, and FFPE samples described with multiplexed PCR-based TCR sequencing using genomic DNA. The T cell structures of the 4 sample types (PBMC, stimulated CD3, LBC, and FFPE) are shown as the total number of T cells defined by nucleotides, productive clonality, fraction of T cells, number of clonotypes defined by nucleotides, maximum productive frequency, and the quantity of DNA (ng) required per cell.

The percentages of the top 15 most frequent clonotypes were significantly increased after 4 vaccinations in all sample types except for FFPE (Figure 3A). Venn diagrams of clonotypes detected in PBMC, LBC, and FFPE at pre-vaccination and post-4 vaccinations revealed that some clonotypes can be detected in all sample types, reflecting the capacity of at least a subset of T cells to traffic to the cervix (Figure 3B). A beta-binomial model, which accounts for variance due to random sampling from a highly diverse repertoire and time-dependent variance for identifying clinically relevant expansion of T cells,^18^ was used to identify putatively vaccine-specific TCRs using pre and post-4 PBMC samples. Seventy putatively vaccine-specific TCRs were identified using the CDR3 nucleotide sequences (Supplementary Table 1). The numbers of such clonotypes and total T cells in pre- and post-4 vaccination FFPE (1 and 9 clonotypes, and 1 and 13 total T cells, respectively) and pre- and post-4 vaccination LBC [14 and 47 clonotypes (Figure 3C), and 33 and 1,523 total T cells respectively] showed that LBC may be more an informative sample type because of greater T cell abundance.

**Figure 3.**
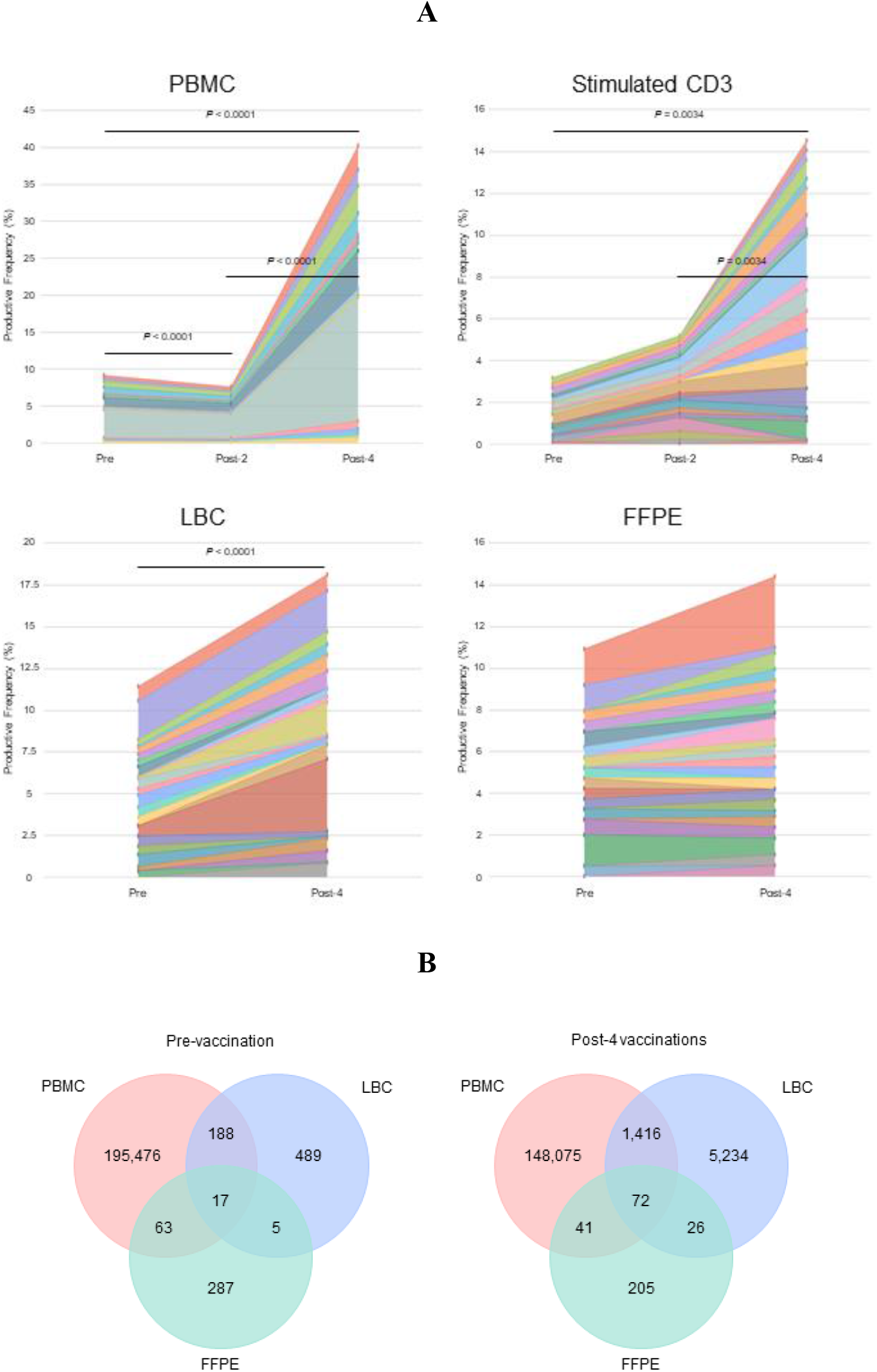
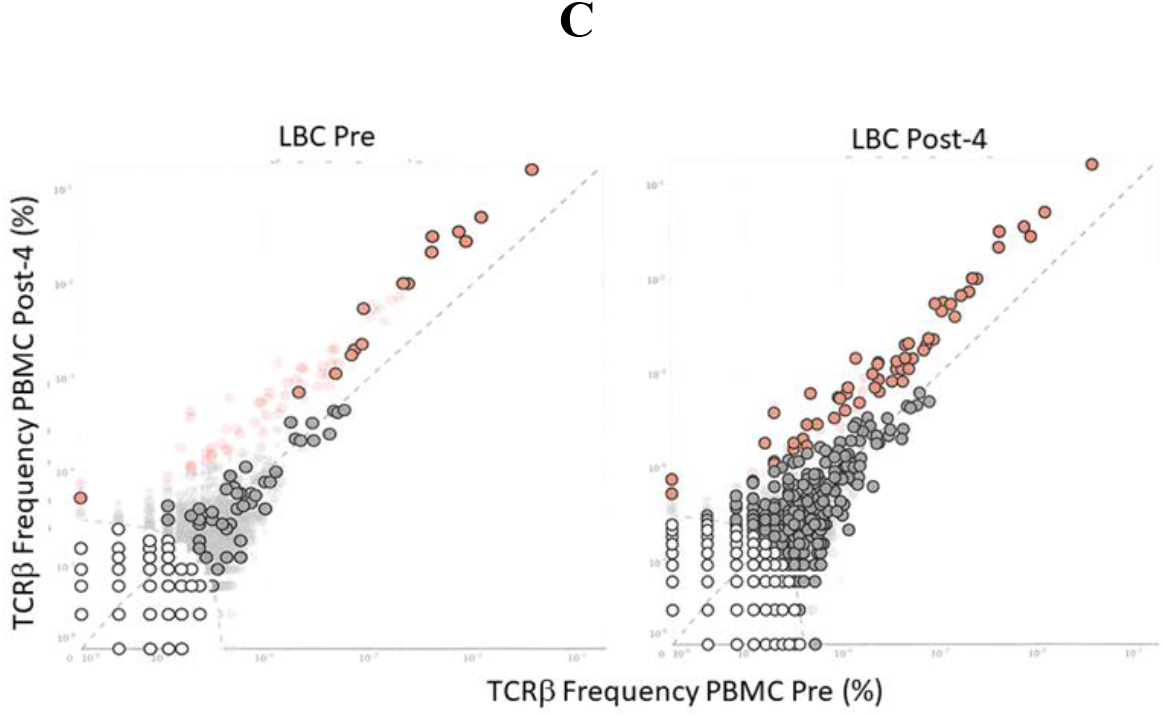
Tracking of clonotypes in the peripheral blood and cervix. **A.** Tracking of the top 15 clonotypes defined by nucleotides are shown in productive frequency. The top 15 highest frequency clonotypes were significantly decreased after 2 vaccinations (Wilcoxon matched-pairs signed-ranks test, *p*=0.0012), but significantly increased after 4 vaccinations (*p*<0.0001) in PBMC samples calculated using the numbers of T cells. For stimulated CD3 T cell samples, significant increases were seen between pre-vaccination and post-4 vaccinations (*p*=0.034) and between post-2 and post-4 vaccinations samples (*p*=0.0034). A significant increase was seen in LBC samples (*p*<0.0001) but not in FFPE samples. **B.** Venn diagrams of clonotypes defined by nucleotides in PBMC, LBC, and FFPE samples pre-vaccination and post-4 vaccinations. Most clonotypes appear only in one sample type, but there are 17 TCRs present at the pre-vaccination visit and 72 TCRs at the post-4 vaccination visit. **C.** Putatively vaccine-specific clonotypes in LBC samples before and after 4 vaccinations. Seventy putatively vaccine-specific clonotypes were identified through a comparison of post-4 PBMC and pre PBMC samples using the beta-binomial model (shown as red dots with and without black circular borders). Red dots with black circular borders represent these putatively vaccine-specific TCRs present in pre-vaccination LBC sample (*n*=15) and in post-4 vaccination LBC sample (*n*=57). Dark grey dots are not significantly different between pre-vaccination and post-4 vaccinations PBMC samples. Dark grey dots with black circular borders are not significantly increased but are present in the respective LBC sample. Light grey dots with black circular borders are not present in the respective LBC sample.

### Single-cell RNA-seq and TCR sequencing

Of 8.5 × 10^6^ peptide (3 15-mer peptides covering the HPV 16 E6 91-115 region)-stimulated and IFN-γ labeled cells, 1.3 × 10^6^ (15.3%) were positively sorted. For the TCR sequencing, the estimated number of cells was 12,240 with mean read pairs of 13,678 per cell. Most (10,246 of 12,240 or 83.7%) cells contained productive V-J spanning pairs. The TCR β amino acid sequences of the 4 clonotypes with a frequency of ≥5% among the IFN-γ positive cells are shown in Table 3.

**Table 3.** TCR β CDR3 sequences of clonotypes with specificity to HPV 16 E6 91-115 and ≥5% frequency.

| Clonotype | Number | Frequency (%) | Amino acid sequence |
| --- | --- | --- | --- |
| 1 | 2,615 | 33.3 | CASSPTSGGLTWDEQYF |
| 2 | 1,340 | 17.0 | CASSHNSGREGNEQFF |
| 3 | 772 | 9.8 | CASSFPGENEQFF |
| 4 | 678 | 8.6 | CASSWEAGQETQYF |

The single-cell RNA-seq analysis revealed an estimated 15,114 total number of cells, 32,659 mean reads per cell, and 2,047 median number of genes per cell. After filtering and normalization, cells were clustered into 9 separated populations (Figure 4A). Notably, abundant expression of IFN-γ and tumor necrosis factor (TNF), but not interleukin-4 (IL-4), was detected in cluster #1, #2 and #3 within the CD8+ T-cell populations, as shown in volcano and feature plots (Figure 4B and 4C).

**Figure 4.**
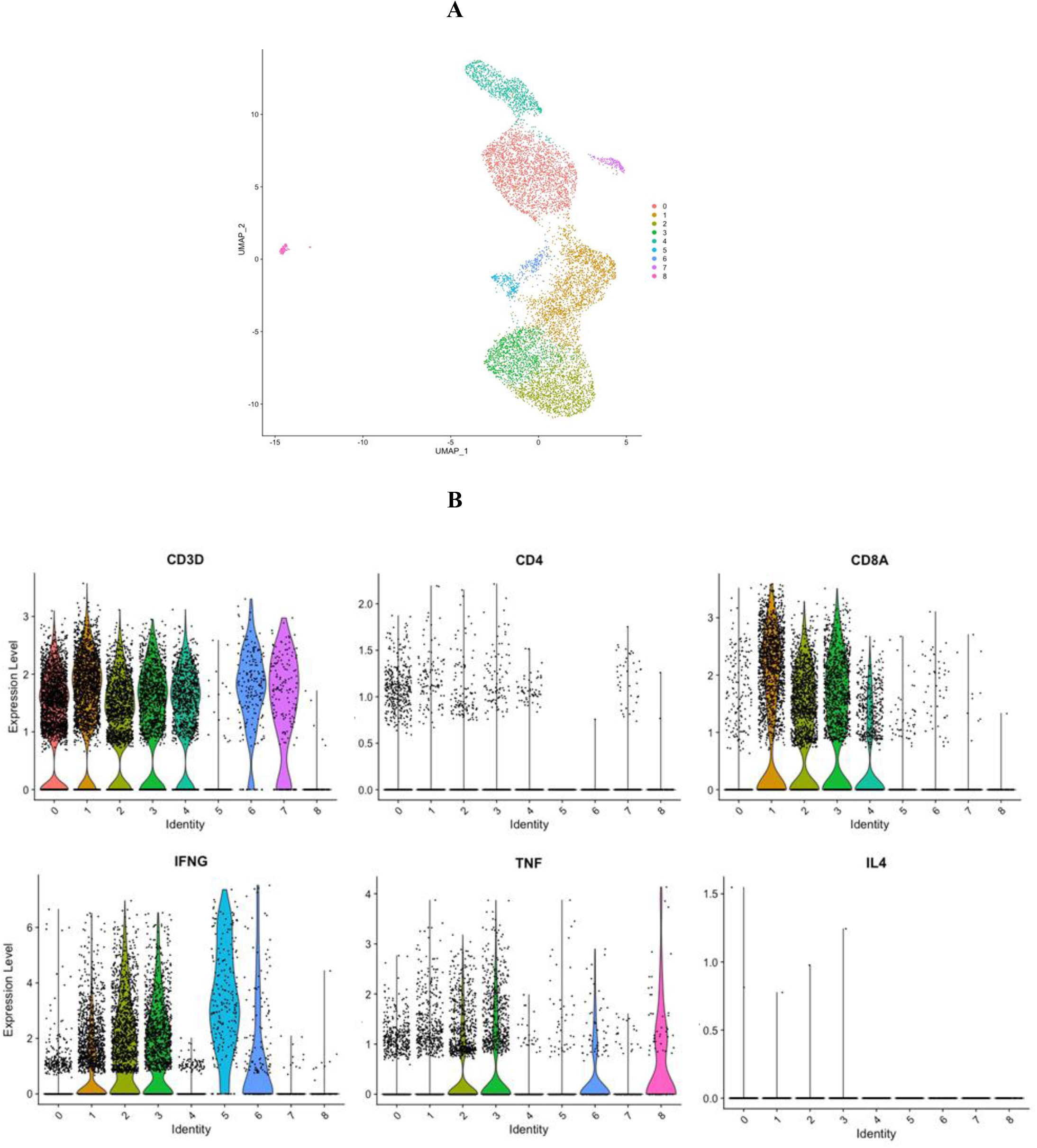
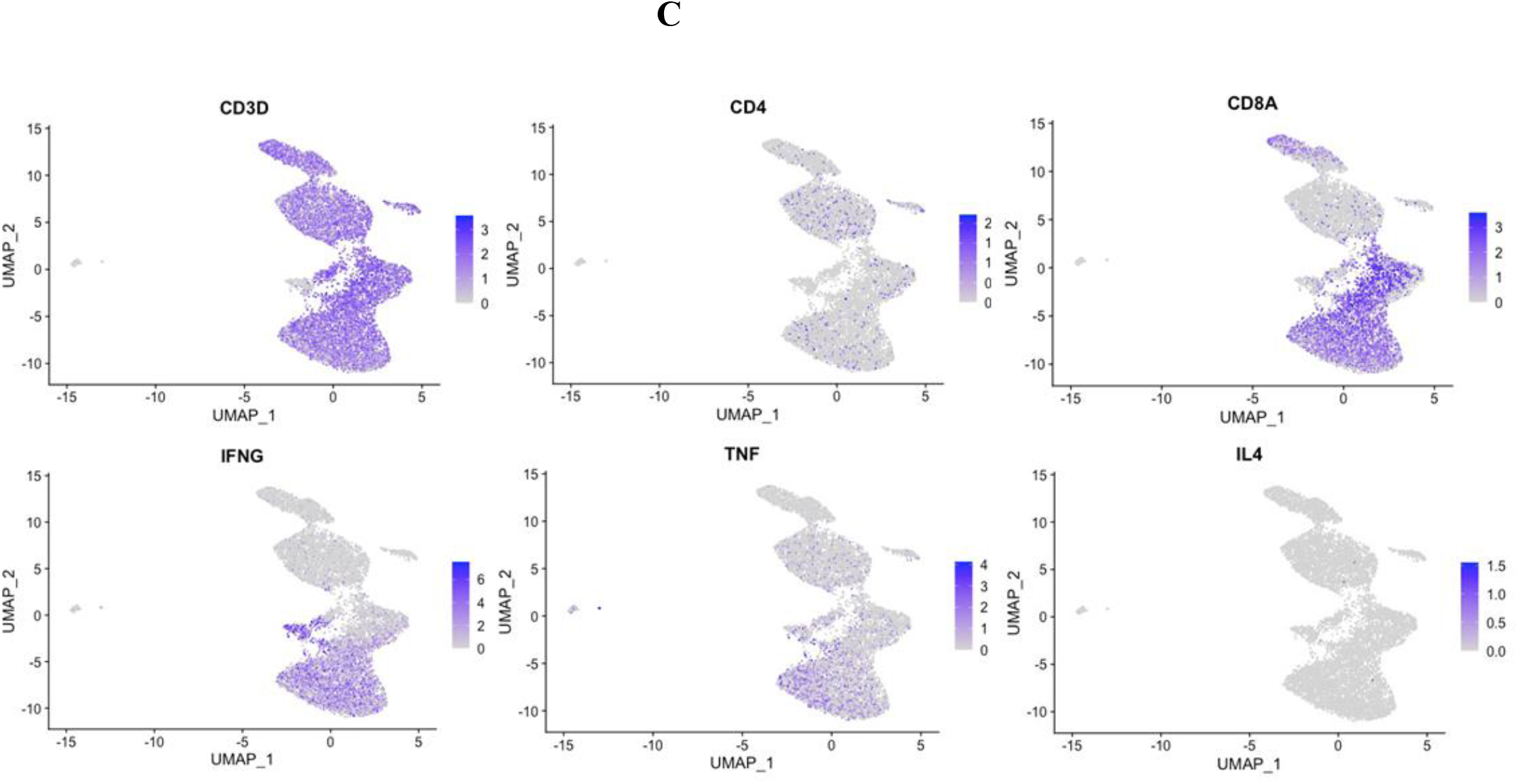
Single-cell gene expression profile of HPV 16 E6 91-115 specific cells. **A.** A UMAP plot showing 9 clusters based on gene expression profiles. **B.** Volcano plots showing CD3D, CD4, CD8A, IFN-γ, TNF, and IL-4 gene expression. **C.** Feature plots showing CD3D, CD4, CD8A, IFN-γ, TNF, and IL-4 gene expression.

### Tracking of the HPV 16 E6 91-115 specific T cells

Using the TCR β CDR3 sequences of the 4 clonotypes specific for HPV 16 E6 91-115, their frequencies in PBMC, LBC, and FFPE samples were determined using TCR β chain sequencing (Figure 5). All 4 clonotypes were detectable in PBMC and LBC prior to vaccination, and their expansion after 4 vaccinations is shown. Only one T cell of clonotype 2 is detectable prior to vaccination in FFPE. All 4 clonotypes were detectable after 4 vaccinations, but only at 2 T cells for clonotypes 1, 3, and 4 and 1T cell for clonotype 2. As much fewer cells were detected in FFPE, LBC was a better source for assessing T cell populations, at least in this subject. All 4 clonotypes were represented in the the top 15 most frequent clonotypes for PBMC, LBC, and stimulated CD3 cells, but only clonotype 1 was present in FFPE (Figure 3A). Of the 70 clonotypes identified to be putatively vaccine-specific using the beta-biomial model, 60 clonotypes were shown to be HPV 16 E6 91-115 specific (Supplementary Table 1). Clonotype 1 was the most abundant clonotype in PBMC and LBC, and the second most abundant clonotype in stimulated CD3.

**Figure 5.**
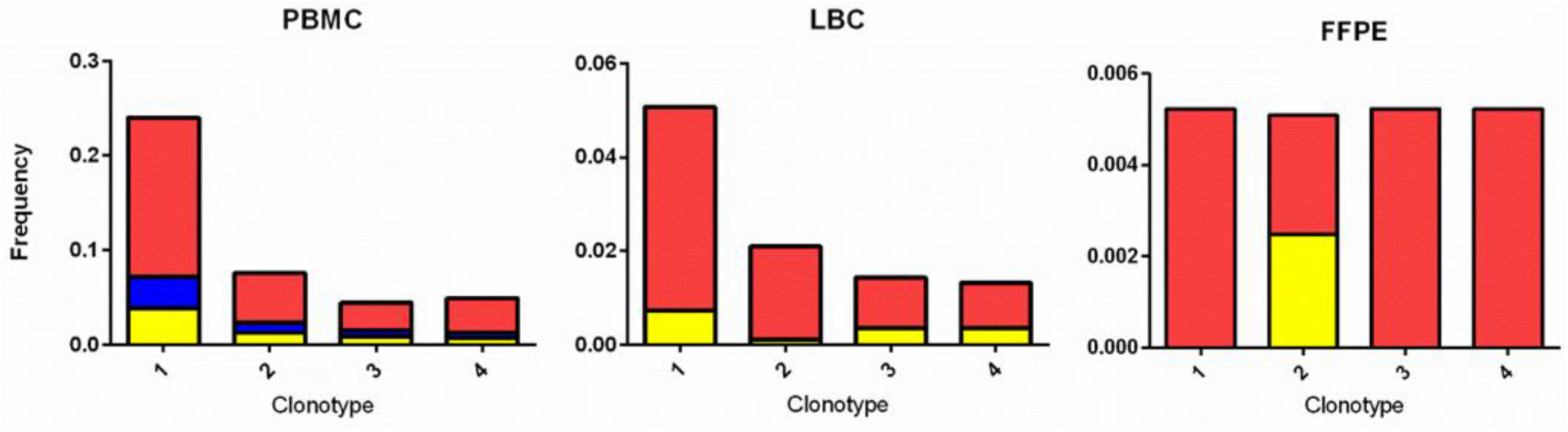
Tracking HPV 16 E6 91-115 specific T cells in PBMC, LBC, and FFPE. The TCR sequences of. The TCR Vα and Vβ sequences of HPV 16 E6 91-115 specific T cells were determined by sorting and sequencing such cells based in IFN-γ secretion upon peptide stimulation. The TCR Vβ CDR3 sequences of top 4 clonotypes (≥ 5% of IFN-γ secreting cells) are shown in Table 3. The frequencies of these clonotypes in PBMC, LBC, and FFPE at prevaccination (yellow), post-2 vaccinations (blue), and post-4 vaccinations (red) time points are shown. All 4 vaccine-specific clonotypes in PBMC and LBC increased in frequency after 4 vaccinations. On the other hand, data from FFPE were not as informative.

## Discussion

This was a proof-of-concept study to demonstrate the utility of TCR analyses using high-throughput sequencing technology in the context of HPV therapeutic vaccine trials. The earliest evidence of the link between HPV and cervical cancer was discovered in 1983 by Harald zur Hausen and his colleagues.^19^ A Nobel Prize was later awarded. To date, over 200 HPV types have been described.^20^ HPV antigens are ideal targets for cancer immunotherapy because they are foreign. Various versions of investigational HPV therapeutic vaccines have been in clinical trials for about the last 30 years, but none has been approved by the United States Food and Drug Administration. Investigational HPV therapeutic vaccines have been tested for many indications including clearance of HPV 16 and/or 18 infection^,21^ HSIL regression,^7,8,22^ prevention of recurrence of squamous cell carcinoma of head and neck (HNC),^23^ treatment of advanced stage cervical cancer,^24,25^ and treatment of advanced stage HNC.^26^ The assessment of vaccine efficacy depends on the indication being tested. For HPV 16 infection clearance, HPV-DNA typing was used,^21^ and biopsies were utilized to evaluate HSIL regression.^7,8,22^ Lack of recurrence within a 2 year period is being used for assessing prevention of recurrence.^23^ Antitumor efficacy was examined using the numbers of patients with complete and partial response, tumor shrinkage, duration of response.^25^

Unlike the HPV prophylactic vaccines which work by inducing production of neutralizing antibodies,^27,28^ the HPV therapeutic vaccines are believed to cast their effects through stimulation of cell-mediated immunity, mainly T cells. Therefore, assessments of T cell immune response should be included in the endpoints of clinical trials. Such implementation varies widely among the clinical trials because T cell assays are technically challenging. In Maciag *et al*.,^24^ the investigators attempted to perform IFN-γ ELISPOT assay using pooled peptides, but most samples were not suitable due to low yield and viability after thawing. Of the 3 patients with a sufficient amounts of cells available to perform the assay, only one demonstrated an HPV-specific T cell response after vaccination. HPV 16 E7 short and long peptides were pooled before testing, so no information as to which portion of the protein contained immunogenic epitopes was obtained.^24^ In the GTL001 trial, van Domme *et al*. performed *ex vivo* IFN-γ ELISPOT assay with pooled HPV 16 E7 peptides or HPV 18 E7 peptides. Overall, 18 of 31 (58.1%) of patients who received any dose of GTL001 with imiquimod demonstrated positive ELISPOT results to either protein. Trimble *et al*. also tested immune responses using IFN-γ ELISPOT assay and intracellular cytokine staining for assessment of T cell immunity. Significantly higher responses were reported for patients who received the VGX-3100 vaccine compared to those who received placebo. As peptides were pooled for each protein tested (HPV 16 E6, HPV 16 E7, HPV 18 E6, and HPV 18 E7), information on which portion of the protein contained the immunogenic epitopes was not determined.^22^ In the clinical trial which treated advance HNC patients with ISA100 and nivolumab, the investigators performed IFN-γ ELISPOT assay for HPV 16 E6 and E7 again using peptide pools. Variable increases in the number of HPV-specific T cells were observed after vaccination in both responders and nonresponders making the role of vaccine-induced T cells uncertain. Furthermore, the immune response did not correlate with efficacy endpoints.^26^ In addition to IFN-γ ELISPOT assay, Melief *et al*. performed lymphocyte stimulation test, intracellular cytokine staining, and cytometric bead arrays to assess immune responses. In all 64 patients who received ISA101 vaccination, HPV 16 E6 and/or E7-specific T cell responses to one or more of 6 peptide pools (4 pools for HPV 16 E6 and 2 pools for HPV 16 7 protein) was demonstrated. Our IFN-γ ELISPOT protocol distinguishes itself among others in that we tested for 10 HPV 16 E6 peptides pools and 6 HPV 16 E7 peptide pools (Fig. 1B).^7,8^ Therefore, the locations of the antigenic epitopes can be narrowed down to 25 amino acid regions. This characteristic of our assay was key to identifying a significant response to the HPV 16 E6 91-115 region, and subsequent isolation of T cells based on IFN-γ secretion. Overall, 61% (19 of 31) of vaccine recipients demonstrated a new T cell response to at least one region of HPV 16 E6 protein.^7,8^ Furthermore, the responses were statistically significant in 42% (13 of 31) of the subjects.^7,8^ However, such responses observed after vaccination in the peripheral blood did not correlate with whether the cervical HSIL lesions regressed. It is possible that the vaccine-induced HPV-specific T cells failed to reach the cervix in some subjects.

TCRs are highly diverse heterodimers consisting of α and β chains in the majority of T cells. However, 1-5% of T cells express γδ chains.^29^ Similar to B cell receptors, the TCR chains contains a variable region responsible for antigen recognition, and a constant region. The variable region of the α and δ chains is encoded by recombined variable (V) and joining (J) genes. Additionally for the β and γ chains, diversity (D) genes are also recombined (i.e., VDJ recombination). Therefore, the β and γ chains are more diverse than the α and δ chains. The advent of high-throughput sequencing made it possible to probe into the complexity of such TCRs. In the current study, we employed TCR β chain deep sequencing using bulk DNA and single-cell RNA-based TCR analysis using mRNA. The former has the advantage of using DNA, which can be extracted from LBC and FFPE samples; therefore, live cells are not necessary. The latter was utilized to analyze IFN-γ secreting HPV 16 E6 91-115 specific T cells. Information on TCR α and β sequences and their parings was obtained, and the gene expression profiles of individual cells was examined. We demonstrated that using the information from a traditional IFN-γ ELISPOT assay in combination with TCR sequencing enables us to demonstrate the expansion of HPV-specific T cells and their presence in the cervix. In addition to demonstrating the information on TCR α and β chain pairings, the single-cell RNA based method has the advantage of yielding the entire sequences of the α and β chains. This would enable construction of the TCRs in viral rectors with which their specificities can be verified.^30,31^ Furthermore, such engineered T cells can be used for immunotherapy as demonstrated by Draper and colleagues^32^. They used T cells genetically engineered to express the TCR of HPV 16 E6 29-38 (TIHDIILECV) epitope restricted by HLA-A*02:01. These engineered T cells were shown to be cytotoxic for HPV 16 positive cervical and HNC cell lines.^32^ The limitation of our current study was that we only examined one subject in this proof-of-concept study. As the Phase II clinical trial of PepCan is ongoing (NCT02481414), additional analyses of Phase II participants would aid in determining the generalizability of the findings of this study.

## Methods

### Ethics

This study was approved by the Institutional Review Board at the University of Arkansas for Medical Sciences (IRB number 130662) and written informed consent was obtained.

### Subject, clinical trial design, and laboratory analyses

This open-label single center dose-escalation Phase I clinical trial of PepCan was reported previously.^7,8^ Subject 6 was selected for the current study because she was a vaccine responder, and sufficient amounts of her samples were available for further analyses. Briefly, subjects qualified for vaccination if biopsy-proven CIN 2 and/or CIN 3 (Figure 1). PepCan (subject 6 received 50 μg/peptide dose) was given 4 times 3 weeks apart, and LEEP was performed 12 weeks after the last vaccination. Cervical LBC samples (ThinPrep, Hologic, Marborough, MA) were collected for HPV typing before vaccination at the time of qualifying biopsy, and after 4 vaccinations at the time of LEEP. Blood was drawn before vaccination, after 2 vaccinations, and after 4 vaccinations, and routine clinical laboratory tests (complete blood count, sodium, potassium, chloride, carbon dioxide, blood urea nitrogen, creatinine, aspartate transaminase, alanine transaminase, lactate dehydrogenase, γ-glutamyl transpeptidase, total bilirubin, and direct bilirubin) were performed. PBMCs were isolated using the ficoll density gradient method. Cells were stored in liquid nitrogen tanks while LBC samples were kept in −80°C freezers. FFPE samples were stored at room temperature.

Research laboratory analyses performed as a part of the clinical trial included HPV typing (Linear Array HPV Genotyping Test, Roche Molecular Diagnostics, Pleasanton, CA), IFN-γ ELISPOT assay, fluorescent-activated cell sorter analysis of peripheral Th1, Th2, and Treg cells, and HLA class I and class II low-resolution typing (One Lambda, West Hills, CA). The Linear Array HPV Genotyping Test detects 37 individual HPV types (6, 11, 16, 18, 26, 31, 33, 35, 39, 40, 42, 45, 51, 52, 53, 54, 55, 56, 58, 59, 61, 62, 64, 66, 67, 68, 69, 70, 71, 72, 73, 81, 82, 83, 84, IS 39, and CP6108). For the ELISPOT assay, magnetically selected CD3 T cells were stimulated with autologous monocyte-derived dendritic cells pulsed with HPV 16 E6 or E7 using recombinant vaccinia viruses and recombinant GST fusion proteins twice with a one-week duration for each stimulation. The assay was performed in triplicates using overlapping HPV 16 E6 and E7 peptides covering HPV 16 E6 1-25, 16-40, 31-55, 45-70, 61-85, 76-100, 91-115, 106-130, 121-145, 136-158 and HPV 16 E7 16-40, 31-55, 46-70, 61-85, and 76-98 regions. PBMCs were stained for CD4, CD25, T-bet, GATA3, and Foxp3. The percentage of CD4 cells positive for T-bet represented Th1 cells, those positive for GATA3 represented Th2 cells, and those positive for CD25 and FoxP3 represented Tregs.

### Multiplexed PCR-based TCR sequencing

The TCR β CDR3 regions were PCR-amplified and sequenced (immunoSEQ, Adaptive Biotechnologies, Seattle, WA)^33^ using genomic DNA from PBMCs (pre-, post-2, and post-4), stimulated CD3 T cells (pre, post-2, and post-4), LBC (pre and post-4), and FFPE (pre and post-4). Using bias-controlled V and J gene primers, the rearranged V(D)J segments were amplified and sequenced. A clustering algorithm was used to correct for sequencing errors, and the CDR3 segments were annotated according to the International ImMunoGeneTicsCollaboration^34,35^ to identify the V, D, and J genes that contributed to each rearrangement. A mixture of synthetic TCR analogs was used in PCR to estimate the number of cells bearing each unique TCR sequence.^36^ “Detailed rearrangements”, “Track Rearrangements”, “Venn Diagram”, “Differential Abundance”, and “Scatterplot with Annotation” features of the immunoSeq analyzer^37^ were used to analyze data.

### Single-cell RNA-seq of HPV-specific T cells

In order to obtain TCR Vα and Vβ sequences of T cells specific for HPV 16 E6 91-115 (Figure 1B), such T cells were selected using a human IFN-γ Secretion Assay – Cell Enrichment and Detection Kit (Miltenyi Biotec, Auburn, CA) following the manufacturer’s instructions as previously described.^38–42^ Post-4 PBMC sample cryopreserved after monocyte depletion was thawed and cultured overnight in Yssel’s media (Gemini Bio Products, West Sacramento, CA) with 1% human serum and 1,200 IU/ml of recombinant human interleukin-2 (R&D Systems, Inc., Minneapolis, MN). As a positive control, healthy donor PBMC mixed with 1% HPV 16 E6 52-61 (FAFRDLCIVY)-specific T cell clone cells were processed in the same manner. The cells were stimulated for 3 h with 10 μM each of peptides in RPMI1640 media plus 5% human serum: FAFRDLCIVY for the positive control, and the three 15-mer overlapping peptides covering the E6 91-115 region (91-105, YGTTLEQQYNKPLCD; 96-110, EQQYNKPLCDLLIRC; 101-115, KPLCDLLIRCINCQK). IFN-γ secreting cells were labeled using the IFN-γ catch reagent and phycoerythrin (PE)-labeled IFN-γ detection antibody. The positive control sample and healthy donor PBMC stained with mouse IgG1K isotype labeled with PE (eBiosciences) was used as a negative control to set the gate. The IFN-γ positive cells were sorted using FACS Aria (BD Biosciences, Franklin Lakes, New Jersey).

A Next GEM Chip G was loaded with approximately 10,000 cells and Chromium Next GEM Single Cell 5’ Library Gel Bead Kit v1.1 reagent (10X Genomics, Pleasanton, CA). An emulsion was generated with the Chromium Controller (10X Genomics). Gene expression (GEX) libraries were prepared with the Chromium Single Cell 5’ Library Construction Kit and TCR libraries were prepared with the Chromium Single Cell V(D)J Enrichment Kit, Human T Cell (10X Genomics). A low-pass surveillance sequencing run of both libraries were performed on separate Illumina mid-output MiniSeq flow cells (GEX library Read1:26bp, Read2:91bp, TCR library Read1:150bp, Read2:150bp). Sequencing was scaled up on an Illumina NextSeq 500 with a high-output 150-cycle v2.5 kit for the GEX library and a mid-output 300-cycle v2.5 kit for the TCR library; both runs used identical read lengths as on the MiniSeq. Data was aggregated from both runs.

Sequencing data were first processed by a Cell Ranger pipeline (v4.0.0; 10X Genomics). Gene expression sequencing data were mapped to human reference (GRCh38-2020A) dataset. The raw single-cell data were processed by R package Seurat v. 3.2.2, by following the recommended steps and settings. The low-quality cells and doublets were filtered out by the following recommended setting: percentage of mitochondrial genes > 5%, number of detected genes < 200 and number of detected genes > 2500. The clustering was performed with the resolution setting at 0.4. The UMAP (Uniform Manifold Approximation and Projection) plot, volcano plots and feature plots were also generated by Seurat (Fig. 4).

TCR sequencing data were mapped to human TCR reference (GRCh38-alts-ensembl-4.0.0) dataset, and they were further analyzed by Loupe V(D)J Browser (v3.0.0; 10X Genomics). T cell clonotypes were defined based on TCR Vβ CDR3 nucleotide sequences after removing single cells containing only α chains and those containing two different TCR Vβ CDR3 nucleotide sequences (likely doublets). For calculating the frequencies of ≥ 5% clonotypes (Table 3), clonotypes with two or more single cells were included. Full-length TCRα/β amino acid sequences were obtained by the Loupe V(D)J Browser.

## Data availability

The bulk TCR Vβ deep sequencing data are available in immuneACCESS (immuneACCESS DOI: https://DOI10.21417/TS2020HPV).

## Statistical analysis

A paired *t*-test was performed to assess the significant changing of spot forming units (i.e., IFN-γ secreting cells) before and after vaccination in ELISPOT assay. The number of T cells between study visits in PBMC, stimulated CD3 T cells, LBC, and FFPE were compared using Wilcoxon matched-pairs signed-ranks test (GraphPad Instat 3, GraphPad Software, San Diego, CA). A *p* value < 0.05 was considered statistically significant.

## Acknowledgements

This work was supported by the grant from the National Institutes of Health (R01CA143130, USA), Drs. Mae and Anderson Nettleship Endowed Chair of Oncologic Pathology (31005156, USA), and the Arkansas Biosciences Institute (the major component of the Tobacco Settlement Proceeds Act of 2000, G1-52249-01, USA).

## Authors’ Contributions

M.N., D.J., To.S. and Y.-C. L developed the concepts and designed this project. Ta.S., S.S., T. E., O. W. S., C.M.Q., H.C. and M.N. performed the experiments. Ta.S., B.L. and M.N. wrote the manuscript, and all authors edited it. Ta.S., S.S., H.J.S., E.P., and M.N. performed statistical and bioinformatics analyses.

